# TRPV4 inhibition prevents increased water diffusion and blood-retina barrier breakdown in the retina of streptozotocin-induced diabetic mice

**DOI:** 10.1101/535526

**Authors:** Maricruz Orduña Ríos, Ramsés Noguez Imm, Nicole Marilú Hernández Godínez, Ana María Bautista Cortes, Wolfgang Liedtke, Ataúlfo Martínez Torres, Luis Concha, Stéphanie Thébault

## Abstract

A better understanding of the molecular and cellular mechanisms involved in retinal hydro-ionic homeostasis imbalance during diabetic macular edema (DME) is needed to gain insights into retinal physio(patho)logy that will help elaborating innovative therapies with lower health care costs. Transient receptor potential cation channel subfamily vanilloid member 4 (TRPV4) plays an intricate role in homeostatic processes that needs to be deciphered in normal and diabetic retina. Based on previous findings showing that TRPV4 antagonists resolve blood-retina barrier (BRB) breakdown in diabetic rats, we evaluated whether TRPV4 channel inhibition prevents and reverts retinal edema in streptozotocin(STZ)-induced diabetic mice. We assessed retinal edema using common metrics, including retinal morphology/thickness (histology) and BRB integrity (albumin-associated tracer), and also by quantifying water mobility through apparent diffusion coefficient (ADC) measures. ADC was measured by diffusion-weighted magnetic resonance imaging (DW-MRI), acquired *ex vivo* at 4 weeks after STZ injection in diabetes and control groups. DWI images were also used to assess retinal thickness. TRPV4 was genetically ablated or pharmacologically inhibited as follows: left eyes were used as vehicle control and right eyes were intravitreally injected with TRPV4-selective antagonist GSK2193874, 24 h before the end of the 4 weeks of diabetes. Histological data show that retinal thickness was similar in nondiabetic and diabetic wt groups but increased in diabetic *Trpv4*^−/−^ mice. In contrast, DWI shows retinal thinning in diabetic wt mice that was absent in diabetic *Trpv4*^−/−^ mice. Disorganized outer nuclear layer was observed in diabetic wt but not in diabetic *Trpv4*^−/−^ retinas. We further demonstrate increased water diffusion and BRB hyperpermeability in diabetic wt mice, effects that were absent in diabetic *Trpv4*^−/−^ mice. Retinas of diabetic mice treated with PBS showed increased water diffusion that was not inhibited by GSK2193874. ADC maps in nondiabetic *Trpv4*^−/−^ mouse retinas showed restricted diffusion. Our data provide evidence that water diffusion is increased in diabetic mouse retinas and that TRPV4 function contributes to retinal hydro-ionic homeostasis and structure under control conditions, and to the development of BRB breakdown and increased water diffusion in the retina under diabetes conditions. A single intravitreous injection of TRPV4 antagonist is however not sufficient to revert these alterations in diabetic mouse retinas.

## Introduction

Diabetic macular edema (DME) is a complication of diabetes that results from an imbalance between retinal fluid entry and fluid exit, leading to intraretinal and subretinal fluid accumulation in the macula region of the retina (Daruich et al., 2018). Under normal conditions, the inner blood-retina barrier (BRB), formed by the tight junctions between the intraretinal vascular endothelial cells, limits fluid entry. Dynamic interactions with pericytes, astrocytes, Müller glia, and microglia also contribute to the inner BRB function. Additionally, retinal fluid entry is limited by the retinal pigment epithelium junctions and the outer limiting membrane (OLM), which both form the outer BRB. Fluid exit is ensured by Müller glia and retinal pigment epithelium that continuously drain water and osmolytes from the retina.

During DME, permeability through retinal barriers increases, causing protein leakage within the interstitial retinal tissue that will be accompanied by water accumulation (Romero-Aroca, 2010). This vasogenic edema may be accompanied by an increase in intracellular fluid volume (cell swelling) (Kohno et al., 1983; Yanoff et al., 1984). Additionally, decreased drainage functions account for decreased fluid exit in DME. Cell loss, OLM disruption, and deregulation of transporters and ion/water channels contribute to BRB breakdown and reduced drainage in DME. These pathological mechanisms are a consequence of fundamental pathways activated by chronic hyperglycemia, including inflammatory ones (Daruich et al., 2018; Klaassen et al., 2013; Willermain et al., 2018).

Recent evidence supports that transient receptor potential cation channel subfamily vanilloid member 4 (TRPV4) participates in drainage functions of the retina. Functionally coupled to acuaporin-4, TRPV4 is necessary for the regulatory volume decrease under hypo-osmotic conditions in retinal Müller glia endfeet (Jo et al., 2015). Furthermore, *Trpv4*^−/−^ mice show disrupted OLM (Arredondo Zamarripa et al., 2017). TRPV4 has also been implicated in the regulation of barrier permeability, but this is subject of debate. While studies showed that TRPV4 activation promotes barrier resistance (Akazawa et al., 2013; Ke et al., 2015), others reported that its stimulation provokes barrier breakdown (Gao and Wang, 2010; Hamanaka et al., 2007; Harteneck and Reiter, 2007; Phuong et al., 2017; Reiter et al., 2006; Suresh et al., 2015; Villalta et al., 2014; Willette et al., 2008; Yin et al., 2016). An additional confounding aspect is that TRPV4 expression is downregulated during diabetes (Hills et al., 2012; Klaassen et al., 2013; Monaghan et al., 2015). Based on inconsistent observations in *Trpv4*^−/−^ mice, it has been suggested that the intricate role of TRPV4 may relate to its presence in different vascular beds and most cellular components of the neurogliovascular unit (Earley et al., 2009; Kim et al., 2016; Redmon et al., 2017). In the retina, TRPV4 is expressed in neurons, Müller glia, astrocytes, endothelial cells, and retinal pigment epithelium (Arredondo Zamarripa et al., 2017; Monaghan et al., 2015; Ryskamp et al., 2011; Zhao et al., 2015).

Reduced TRPV4 expression may be a compensatory effect in diabetes. Indeed, TRPV4 activation mediates and amplifies inflammatory responses (Alessandri-Haber et al., 2006; Alessandri-Haber et al., 2004; Alessandri-Haber et al., 2005; Alessandri-Haber et al., 2003; Matsumoto et al., 2018; Okada et al., 2016; Reiter et al., 2006; Vergnolle et al., 2010), including edema (Reiter et al., 2006; Vergnolle et al., 2010). TRPV4 blockade resolves edema in several organs (Balakrishna et al., 2014; Cheung et al., 2017; Jie et al., 2015; Lu et al., 2017; Thorneloe et al., 2012; Zhao et al., 2018) and TRPV4 selective antagonists (RN-1734 and GSK2193874) mitigate BRB breakdown in diabetic rats (Arredondo Zamarripa et al., 2017). The role of TRPV4 in retinal hydro-ionic homeostasis should therefore be better characterized under control and diabetic conditions.

We addressed this issue by first analyzing retinas of *Trpv4*^−/−^ mice subjected to 4 weeks of experimental diabetes induced by streptozotocin. Retinal edema was assessed by measuring morphology/thickness, BRB integrity, and water diffusion using histology, an albumin-associated tracer, and diffusion-weighted magnetic resonance imaging (DW-MRI), respectively. DW-MRI is a MRI modality that can measure apparent diffusion coefficient (ADC), a sensitive biomarker of water mobility (Ebisu et al., 1993), whose increase is observed in models of vasogenic edema (Sood et al., 2009). A previous study showed diffuse retinal edema in diabetic rats using *in vivo* DWI (Berkowitz et al., 2012). MRI analysis showed that diabetic male C57BL/6 mouse retinas are not thicker than their controls (Berkowitz et al., 2009), but retinal water content and diffusion remain to be measured to determine whether or not this model displays retinal edema. Even though mice have been used less frequently than rats as models in studies of DME, they exhibit features of diabetic retinopathy (Hammes et al., 2002; Martin et al., 2004). Then, we examined the retinal outcome after a single intravitreal injection of a very potent and selective TRPV4 antagonist, GSK2193874 (Thorneloe et al., 2012) in diabetic mice.

## Methods

### Reagents

The TRPV4 antagonist GSK2193874 and all other reagents were purchased from Sigma-Aldrich (St Louis, MO).

### Ethics statement

All experiments were approved by the Bioethics Committee of the Institute of Neurobiology at the National Autonomous University of Mexico (UNAM, protocol #74), and methods were carried out in accordance with the National Institutes of Health Guide for the Care and Use of Laboratory Animals, the ARVO Statement for the Use of Animals in Ophthalmic and Vision Research, and with authorization from the Institutional Animal and Care Use Committee.

### Animals

C57BL/6J mice of either sex (5-7 weeks old) were obtained from commercial suppliers, whereas *Trpv4*^−/−^ mice (Liedtke and Friedman, 2003) were a kind gift from Dr. Wolfgang Liedtke (Duke University). Animals were fed *ad libitum* and reared in normal cyclic light conditions (12 h light: 12 h dark) with an ambient light level of approximately 400 lux.

Diabetes was induced with intraperitoneal injections of streptozotocin (60 mg/kg) once a day for five consecutive days (Navaratna et al., 2007). Animals with glucose levels greater than 250 mg/dL after a 6-h fast (Han et al., 2008) were used 4 weeks after diabetes induction. Nondiabetic groups received intraperitoneal injections of citrate buffer once a day for five consecutive days (controls). Body weight and glycemia were monitored weekly (Supplemental Fig. 1).

In addition to the wild-type and *Trpv4*^−/−^ mice rendered or not diabetics, the study included 4 additional groups; control and diabetic mice that were randomized to receive an intravitreal injection with PBS or TRPV4 antagonist. The final injection volume was 0.5 μl. In both diabetic and nondiabetic mice, the right eye was injected with vehicle (PBS) and the left eye received GSK2193874 (17.3 pg, corresponding to 50 nM, as the estimated volume of mouse vitreous is 4.4 μl (Kaplan et al., 2010)). We previously demonstrated that PBS is an acceptable vehicle (Arredondo Zamarripa et al., 2017). Intravitreal injection was performed 24 hours before the end of the 4 weeks of diabetes to mimic a treatment scheme.

Mice were anesthetized intraperitoneally with 70 % ketamine and 30 % xylazine (1 ml/g body weight) before intravitreal injections and MRI procedures. To maintain light conditions between all groups, and given that the magnet bore was dark, mice were manipulated under dim red light and imaging was performed in dark-adapted mice (for at least 12 h).

### Ex vivo MRI procedures

Anesthetized mice were perfusion-fixed with 4 % paraformaldehyde and gadolinium (2 μM) in PBS and stored at 4 °C (D’Arceuil et al., 2007). Mice were decapitated after fixation. Samples were allowed to stabilize at room temperature (21 ± 1 °C) for 4 h before image acquisition. High-resolution anatomic and ADC data were acquired using a 7.0 T system (Bruker Pharmascan 70/16; Billerica, MA, USA), equipped with a gradient set with Gmax=760 mT/m. To enhance signal-to-noise ratio, we used a two-channel Helium-cooled phased-array surface probe (Cryoprobe, Bruker), centered between both eyes. An off-resonance (i.e., B0) map was obtained and used to calculate high-order shim gradients through routines provided by the manufacturer (i.e., MapShim). Images were acquired using a spinecho sequence with three-dimensional spatial encoding, TR = 1000 ms, TE = 21.55 ms, FOV = 12 x 9.04 x 2.4, matrix dimensions = 266 x 200 x 8, yielding a voxel resolution of 45 x 45 x 300 μm^3^, bandwidth = 30.864 kHz, NEX = 1. Slices were oriented perpendicular to the rostro-caudal axis, with imaging planes covering both eyes. Spectral fat suppression was performed using a preparation pulse with bandwidth = 1050 Hz. DWI were obtained with three orthogonal diffusion encoding directions with b = 1200 s/mm^2^, Δ = 8.5 ms, δ = 2.5 ms. In addition, two non-diffusion weighted volumes (i.e., b=0 s/mm^2^) were obtained with identical parameters. Total data acquisition time was 1 h 40 min. Experiments were performed at room temperature controlled at 21 ± 1 °C. ADC maps were calculated as ADC = (ln(S/S_0_)) / −b, where S is the mean of the three DWI and S_0_ is the mean of the two non-diffusion weighted volumes.

### MRI data analysis

Images were analyzed using ITK-SNAP (Yushkevich et al., 2006). As discussed in [36], we inferred layer locations based on the retina’s well-defined laminar structure and clear anatomical landmarks like the vitreous-retina and neuroretina-choroid/retinal pigment epithelium borders. For thickness and ADC analyses, the retina was radially subdivided into eight contiguous segments with 20° angle increments (see Supplemental Fig. 2). The average of the three DWI were used to assess retinal thickness, since these images allow for clear visualization of the retina as hyper-intense band. Manual segmentation of the retina was performed, and average ADC values within the segmented regions were obtained for each radial segment.

Non-diffusion weighted images were used to check retinal structure after intravitreal injections. Of note, anatomical MRI revealed that some *Trpv4*^−/−^ eyes (2 out of 11) showed an enlarged lens capsule (Supplemental Fig. 3).

### Histology

Mouse eyes were fixed for 3 hours at room temperature in 4 % paraformaldehyde. Eyes were then cryopreserved at 4 °C for 1 h 30 in 10 % and 20 % sucrose, respectively, in 30 % sucrose overnight, embedded in Tissue-Tek and frozen with liquid nitrogen. Cryostat was sectioned at 14 μm and mounted on slides for hematoxylin and eosin staining (Arredondo Zamarripa et al., 2017). Retinal layer thickness was quantified, as previously described (Arredondo Zamarripa et al., 2017).

### Statistical analysis

All results were replicated in three or more independent experiments. All data were reported as mean ± s.e.m. All data showed normal distribution and equal variance according to D’Agostino-Pearson omnibus and Levene’s tests, respectively. Comparisons between groups were determined by ANOVA (R Studio). Differences between means with *P* < 0.05 were considered statistically significant.

## Results

### Group summary

*Trpv4*^−/−^ mice had body weight and glycemia comparable to those of wt mice (18.6 ± 0.6 g; *n* = 8 vs. 19.4 ± 1.1 g; *n* = 10 and 125.3 ± 8.4 mg/dl; *n* = 8 vs. 199.3 ± 7.8 mg/dl; *n* = 10, respectively; *P* > 0.05; supplemental Fig. 1). The 4-week streptozotocin treatment did not alter the body weight of wt and *Trpv4*^−/−^ mice (19.4 ± 0.4 g; *n* = 11 vs. 18.3 ± 0.5 g; *n* = 9; *P* > 0.05), but it induced hyperglycemia at similar levels in both groups (378.4 ± 38.8 mg/dl; *n* = 11 vs. 324.7 ± 4.7 mg/dl; *n* = 9, *P* < 0.001, supplemental Fig. 1). Intravitreal injections of PBS or GSK2193874 did not modify nondiabetic and diabetic mouse body weight and glycemia (*P* > 0.05, data not shown).

### TRPV4 contributes to retinal structure and is necessary for BRB rupture and increased water diffusion in diabetic mouse retinas

The histology of *Trpv4*^−/−^ retinas appeared relatively similar to that of wild-type mice except for the OLM that is blurry and the spaces between photoreceptor nuclei that are reduced in *Trpv4*^−/−^ retinas (Fig. 1A). Cell bodies can also be seen in both the outer and inner plexiform layers of *Trpv4*^−/−^ retinas (Fig. 1A). Compared with nondiabetic wt retinas, diabetic wt retinas showed a disorganized outer nuclear layer with photoreceptor cell bodies that lost their round shape and alignment and were separated by large irregular spaces. Additionally, the OLM was not visible, cell bodies could be detected in the outer plexiform layer, and the ganglion cell bodies became oval in diabetic wt retinas (Fig. 1A). The histology of diabetic *Trpv4*^−/−^ retinas appeared similar to that of nondiabetic *Trpv4*^−/−^ retinas; spaces between photoreceptor nuclei looked bigger and the inner nuclear layer thicker (Fig. 1A). Compared to diabetic wt retinas, diabetic *Trpv4*^−/−^ retinas displayed an organized outer nuclear layer and thicker inner retina (Fig. 1A). Total and inner retina thickness was increased in diabetic *Trpv4*^−/−^ retinas (Fig. 1B). We found no significant changes in retinal pigment epithelium (not shown) and outer retinal layer thickness (Fig. 1B).

**Figure 1.**
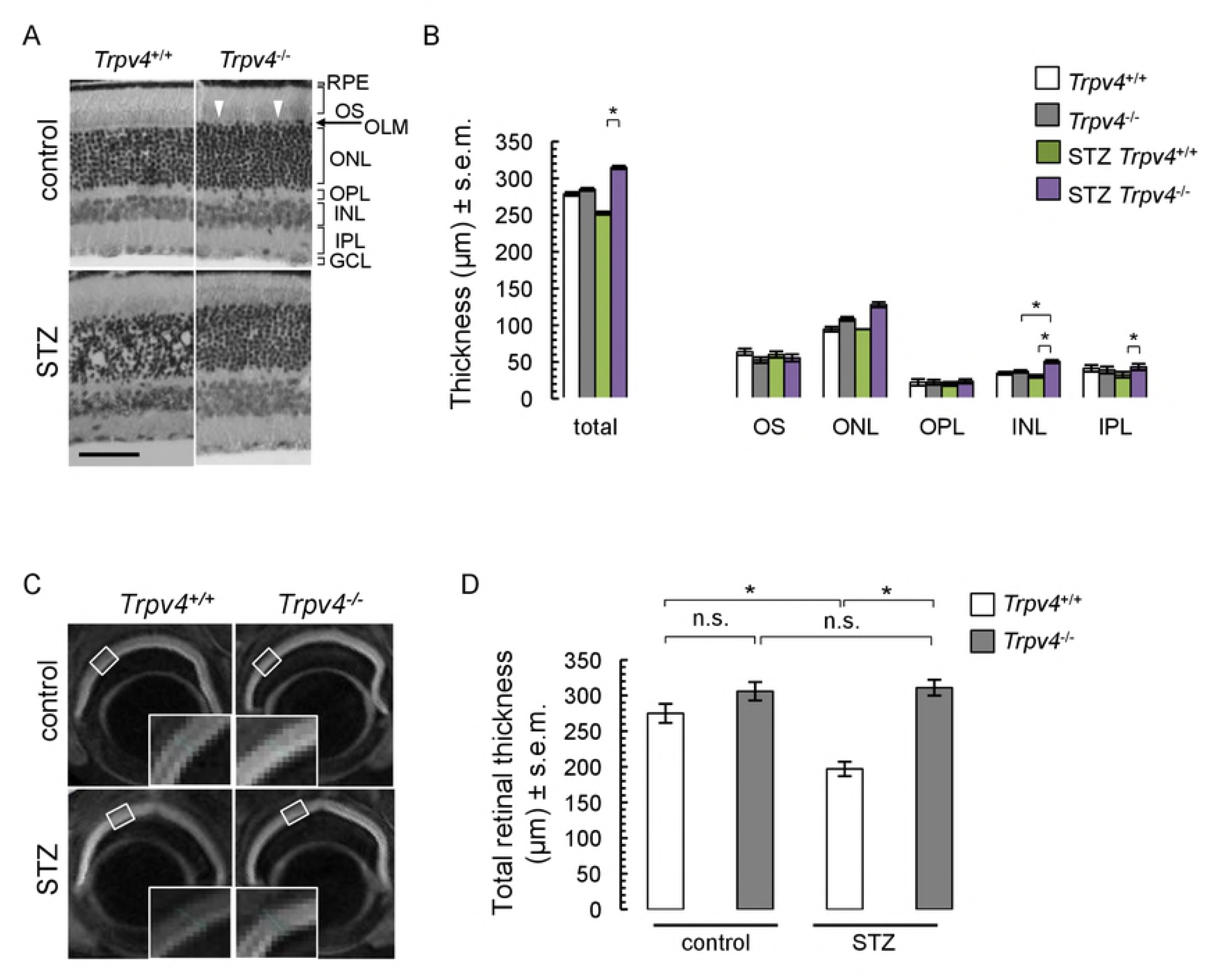
Retinal thickness analysis in nondiabetic (control) and diabetic (STZ) wild-type (*Trpv4*^+/+^) and *Trpv4*^−/−^ mice. Representative images of hematoxylin/eosin-stained retinas (**A**) and corresponding quantification of retinal thickness (**B**). Retinal pigment epithelium (RPE), outer segments (OS), outer nuclear layer (ONL), outer limiting membrane (OLM), outer plexiform layer (OPL), inner nuclear layer (INL), inner plexiform layer (IPL), and ganglion cell layer (GCL). White headarrows indicate disturbed OLM in *Trpv4*^−/−^ retinas. Three sections per retina from each of six animals per group were analyzed. (**C**) Representative DWI images with enlargements corresponding to white squares in control and STZ *Trpv4*^+/+^ and *Trpv4*^−/−^ mouse eyes. (**D**) Summary of retinal thickness measured in DWI images in three separate groups of control and STZ *Trpv4*^+/+^ (*white*) and *Trpv4*^−/−^ (*grey*) mice (μm, mean ± s.e.m.; n = 8-11). *, significant differences of P < 0.05; n.s., not significant.

The retina was easily observable through DWI in all groups (Fig. 1C) and appeared thinner in the streptozotocin-treated wt animals (Fig. 1D). Quantification of DWI-retinal thickness showed that diabetic wt mice had thinner retinas compared to nondiabetic wt mice (Fig. 1E). This effect was absent in diabetic *Trpv4*^−/−^ mice (Fig. 1E). Retinal thickness in nondiabetic *Trpv4*^−/−^ mice was indistinguishable from that of nondiabetic wt mice (Fig. 1E). These effects were not focal but observed throughout the retina (not shown).

Quantification of ADC maps (Fig. 2A) showed an increase (+ 25 ± 2 %) in retinal mean ADC values in diabetic wt mice (Fig. 2B), an effect that was absent in diabetic *Trpv4*^−/−^ mice (Fig. 2B). Nondiabetic *Trpv4*^−/−^ retinas displayed a 20 ± 1 % decrease in mean ADC values compared with the nondiabetic wt group (Fig. 2B). To support these observations, we measured BRB breakdown using the Evans blue technique, given that in the streptozotocin mouse preclinical model of diabetes, BRB breakdown has been shown to occur as early as 2 weeks and up to 24 weeks post-streptozotocin treatment (Hossain et al., 2016; Kim et al., 2012; Leal et al., 2007; Li et al., 2017; Zhou et al., 2014). We confirmed the BRB breakdown in diabetic mice by showing that retinal accumulation of Evans blue-stained albumin tripled (Fig. 2C). The streptozotocin-induced BRB breakdown was not observed in *Trpv4*^−/−^ mice (Fig. 2C). Lack of *Trpv4* did not modify the basal transport through the BRB (Fig. 2C).

### Effect of single intravitreal injection of TRPV4 antagonist GSK2193874 in diabetic mouse retina

Slight retinal/vitreous interface disruption at the intravitreal injection site was observed on the non-diffusion weighted images, but general structure was overall well conserved (Fig. 3A). DWI data (Fig. 3B) demonstrated that injection of either vehicle or GSK2193874 did not affect total retinal thickness (Fig. 1D vs. 3C). Diabetic wt mice showed thinning of the retina if vehicle was administered (Fig. 3C). This overall effect was not prevented by TRPV4 antagonist administration (Fig. 3C). Retinal thinning occurred throughout the entire retina of diabetic mice and GSK2193874 rescued retinas from streptozotocin-induced thinning 0 to 50° from the optic nerve (nasal part of the retina) (Fig. 3D). Using ADC maps (Fig. 4A), we found a streptozotocin-induced increase in mean ADC values (Fig. 4B) that was still present in the eyes of diabetic mice treated with GSK2193874 (Fig. 4B). GSK2193874 did not modify mean ADC values in nondiabetic mice (Fig. 4B). Similar to Figure 3D, Figure 4C shows that mean ADC values increased throughout the entire retina of diabetic mice and that GSK2193874 did not block this increase.

## Discussion

This study examines whether endogenous TRPV4 regulates retinal hydro-ionic homeostasis under normal and diabetic conditions, using common metrics of retinal edema (morphology/thickness and BRB integrity) and DW-MRI. First, we observed that histological and MRI measurements of retinal thickness agreed in nondiabetic mice but not in diabetic mice. Retinal morphometry changes that include OLM disruption and outer nuclear layer disorganization with larger internuclear spaces coincide with BRB breakdown and increased ADC in diabetic mouse retinas. Genetic deletion of *Trpv4* prevents these changes. Lack of TRPV4 induces *per se* qualitative and quantitative changes in the retina, including OLM disruption and restricted water diffusion. Furthermore, we found that a single administration of the selective TRPV4 antagonist GSK2193874 had no effect of retinal thickness and water diffusion in nondiabetic and diabetic retinas.

Our previous work showed that TRPV4 antagonists have significant therapeutic potential for the control of BRB breakdown in DME since they inhibit excessive permeation in the retina of the streptozotocin preclinical rat model of diabetes (Arredondo Zamarripa et al., 2017). Measurement of BRB integrity is however insufficient to prove edema (Bellhorn, 1984; Stefansson et al., 1987). MRI-based analysis of co-localized measures of retinal thickness, BRB breakdown, and water content and mobility has been established as a complete approach to quantify retinal edema in diabetic rats (Berkowitz et al., 2012). We used mice because of the knockout mouse technology to block TRPV4 and retinal water content was not assessed because its quantification using MRI is based on assumptions.

The present data show the retina as a single MR-detected layer, as previously reported (Berkowitz et al., 2004; Roberts et al., 2003). Our MRI estimation of total retinal thickness in control wt mice (275.5 ± 6.1 μm) compares reasonably well with our own values using histology (278.7 ± 2.6 μm) and also with published values using histology, anatomical MRI and OCT from the same mouse strain (270.4 ± 2.3 μm (Cammalleri et al., 2017), 237 ± 3 μm (Berkowitz et al., 2015) and 222.3 ± 2.1 μm (Dysli et al., 2015), respectively). This, however, does not stand true in diabetic mice, where MRI-based measurements showed retinal thinning and histological ones did not. Additionally, a previous MRI study showed that retinal thickness was similar between control and diabetic male C57BL/6 (Berkowitz et al., 2009). In addition to possible variations between C57BL/6 mouse sub-strains, some critical differences (e.g. 1 month vs. 3 months of diabetes and *ex vivo* vs. *in vivo* imaging conditions, current and (Berkowitz et al., 2009) studies) may account for this inconsistency. Because retinal fluid handling is not conserved in diabetic retinas (Berkowitz et al., 2012; Pannicke et al., 2006), the discrepancy between our MRI and histological values may also be attributed to differences in the dehydration process between eyes that were dissected prior PFA fixation in histology procedures, but not in MRI. Furthermore, histological examination has produced inconsistent results with thicknesses that are unchanged (Ganapathy et al., 2009; Martin et al., 2004) or reduced (Ren et al., 2017) at 4-5 weeks after onset of diabetes. At longer periods of diabetes, retinal thinning (Li et al., 2010; Martin et al., 2004; Ozaki et al., 2018; Qian et al., 2018; Sachdeva et al., 2018; Yang et al., 2018; Zheng et al., 2007) or thickening (Berkowitz et al., 2009; Li et al., 2013) has been observed. Retinal thinning concurs with cell loss (Martin et al., 2004; Nishimura and Kuriyama, 1985) and not water loss (Berkowitz et al., 2012) in diverse rodent models of short-term diabetes (< 4 months). Diffuse central retinal edema causes retinal thickening in rats (Berkowitz et al., 2012). Current evidence shows that indicators of retinal edema, i.e. BRB breakdown and increased water diffusion, do not necessarily associate with retinal thickening, if they match morphometric changes compatible with retinal edema (e.g. large internuclear spaces). Histological examination validates diffusion MRI.

If no vasodilation occurs, increased water diffusion in a tissue corresponds to edema. Vasodilation has not been reported in retinas of diabetic animals (Elms et al., 2013; Kaneko et al., 2006; Nakazawa et al., 2008) or of patients with early stages of diabetic retinopathy (Clermont et al., 1997; Grunwald et al., 1996). Our data therefore indicate that diabetic male C57BL/6 mice develop retinal edema. However, retinal water content remains to be estimated in diabetic C57BL/6 mice to undoubtedly conclude that this model displays retinal edema. Edema can be cytotoxic (intracellular) or vasogenic (extracellular); both occur in diabetic retinopathy (Romero-Aroca et al., 2016). Supernormal ADCs are usually found in models of BRB breakdown presumably because free water contained in leaking fluid accumulates into the extracellular space (Berkowitz et al., 2012; Lu and Lei, 2014; Sood et al., 2009). In contrast, the intra-cellular environment is thought to restrict water diffusion, rendering low ADC values in the case of cytotoxic edema. Our results likely identify vasogenic edema in diabetic mouse retinas because they show increased water diffusion and loss of BRB integrity. Nevertheless, we cannot specifically attribute the large irregular spaces between photoreceptor nuclei to increased extracellular space or intracellular cell swelling.

The reason for subnormal ADC values in *Trpv4*^−/−^ retinas is somewhat unclear. Intracellular swelling may cause reduced water mobility. In this sense, TRPV4 mediates regulatory volume decrease - i.e., decrease in swelling in the presence of sustained hypotonic stress (Hoshi et al., 2018; Willermain et al., 2018)- by transducing increases in Müller cell (Jo et al., 2015) and retinal neuron (Ryskamp et al., 2014) volumes into Ca^2+^ signals. However, hematosin-eosin staining suggests greater packing density in *Trpv4*^−/−^ retinas compared to wild-type ones. Elimination of TRPV4 may therefore promote aqueous drainage from the retina. Together with aquaporin-4 and Kir4.1 channels, TRPV4 forms a functional complex that maintains the steady-state ‘osmo-tensile’ homeostasis at Müller glial endfeet (Iuso and Krizaj, 2016; Jo et al., 2015). More work is needed to test whether TRPV4 contributes to retinal hydro-ionic homeostasis through an extended vascular-glial-RPE-ganglion cell network, as shown in other sensory systems (Liedtke and Friedman, 2003; Liedtke and Kim, 2005; Liedtke et al., 2003). Additionally, subnormal ADC can also relate to vasoconstriction and literature supports that TRPV4 activation causes vasodilation (Kohler et al., 2006; Willette et al., 2008).

We found that TRPV4 deletion abolishes retinal edema in streptozotocin-treated mice, demonstrating that TRPV4 contributes to the formation of retinal edema in diabetic conditions. Our finding coincides with the promoting effect of TRPV4 agonism on barrier permeability (Hamanaka et al., 2007; Harteneck and Reiter, 2007; Reiter et al., 2006; Vergnolle et al., 2010; Villalta et al., 2014; Yin et al., 2016) 65]. That streptozotocin-treated mice likely present vasogenic edema (present data) and that this latter associates with inner BRB breakdown (Klaassen et al., 2013), suggests that TRPV4 act on the inner BRB components. In apparent contrast with our data, expression levels of TRPV4 decrease in retinal microvascular vessels from streptozotocin-induced diabetic rats (Arredondo Zamarripa et al., 2017). Nevertheless, TRPV4 is still functional in the diabetic retina (Arredondo Zamarripa et al., 2017; Monaghan et al., 2015). The activity of remaining TRPV4 may be excessive since the amount and nature of TRPV4 endogenous agonists and the levels of glycosylation, free intracellular Ca^2+^, and membrane cholesterol, all of which regulate TRPV4 activity, are altered in diabetic milieu (Arruda and Hotamisligil, 2015; Cohen et al., 2013; Lakk et al., 2017; Xu et al., 2006). As previously proposed (Arredondo Zamarripa et al., 2017), reduced TRPV4 expression may be a compensatory effect in diabetes.

Our data further suggest that TRPV4 participates in the effect of diabetes on retinal fluid accumulation. Based on the facts that (i) VEGF-mediated angiogenesis and inflammation act interdependently in the development of DME (Romero-Aroca et al., 2016), (ii) TRPV4 agonism associates with increased expression of VEGF (Chen et al., 2018) and amplifies inflammatory cascades (Alessandri-Haber et al., 2004; Alessandri-Haber et al., 2005; Alessandri-Haber et al., 2003; Matsumoto et al., 2018; Vergnolle et al., 2010), and (iii) TRPV4 antagonism and anti-angiogenic molecules synergize by activating complementary pathways to counteract the diabetes-like effects on outer BRB permeability (Arredondo Zamarripa et al., 2017), it is possible that TRPV4 blockade targets both the vasogenic and inflammatory pathways in diabetic retina (Pairet et al., 2018). *Trpv4*^−/−^ mice treated with streptozotocin are hyperglycemic, in contrast to littermates subjected to a high-fat diet (Ye et al., 2012), suggesting that the protective effect of TRPV4 inhibition against retinal edema is independent from its protective effect against insulin resistance. The exact underlying mechanism needs to be clarified, but the fact that TRPV4 deletion prevents retinal edema formation associated with diabetes argues in favor of TRPV4 as an etiological agent for DME.

GSK2193874 has no reported systemic side effects (Thorneloe et al., 2012; Vincent and Duncton, 2011) and we confirm herein that short-term inhibition of TRPV4 does not alter BRB permeability (Arredondo Zamarripa et al., 2017). Previous findings showed that TRPV4 blockade preserves the lung (Thorneloe et al., 2012) and brain from vasogenic edema (Zhao et al., 2018). We found that a single injection of GSK2193874 does not revert streptozotocin-induced retinal edema in mice but locally mitigates retinal thinning. The effect appears limited to a peripheral region deviated towards the temporal side, corresponding to the injection site. We consider that a scheme of repeated injections of GSK2193874, as well as the generation of ophtalmological presentations for TRPV4 antagonists may improve this current limitation. While delivery of GSK2193874 has some injection-related side effects (Quiram et al., 2006; Sakamoto et al., 2011), they are uncommon and the intravitreal route ensures that the drug reaches the target site. In this line, we emphasize that the availability of potent and selective drugs to inhibit TRPV4 is very important, not only because all 26 members of the mammalian TRP family are present in the retina (Gilliam and Wensel, 2011), but also to favor the curative use of GSK2193874. Because type 1 and type 2 models of diabetes exhibit similar early signs of diabetic retinopathy (Grunwald et al., 1996), our findings may contribute to treatment development for DME arising from both type 1 and type 2 diabetes. This is of importance considering that more than 90 % of diagnosed patients suffer from a type-2 diabetes phenotype. Further studies using *in vivo* DWI and OCT are also warranted to help determine responses to potential treatment.

We conclude that TRPV4 contributes to the cascade of events that involve the breaking of BRB, which is the stage prior to DME onset. Further experiments are needed to test whether selective inhibition of TRPV4 by GSK2193874 has curative potential for patients with DME. The role of TRPV4 may also extend to other types of retinal edema.

## Acknowledgements

R. Noguez Imm is a Doctoral student from the Programa de Posgrado en Ciencias, Universidad Nacional Autónoma de México (UNAM) and received fellowships from the National Council of Science and Technology of Mexico (CONACYT). We thank E. Espino, M. Ramírez Romero, M. García, A. Castilla, and J.J. Ortiz for their technical assistance, and J. Norris for critically editing the manuscript. This study was supported by the UNAM grant IN209317 (ST).

## Author contributions

Conceived and designed the experiments: ST and LC. Performed the experiments: MOR, RNI, NMHG, ANBC, LC, and ST. Analyzed the data: MOR, LC, and ST. Interpreted the data: WL, LC, AMT, and ST. Contributed reagents/materials/analysis tools: LC, AMT, WL, and ST. Wrote the paper: ST. Critical revision for intellectual content: LC, AMT, WL, and ST. All authors finally approved the submitted version.

## Figure legends

**Figure 2**. Estimated water diffusion and BRB permeability in retinas of control and STZ *Trpv4*^+/+^ and *Trpv4*^−/−^ mice. (**A**) Representative images of ADC maps in control and STZ *Trpv4*^+/+^ and *Trpv4*^−/−^ mouse eyes. (**B**) Summary of ADC values in three separate groups of control and STZ *Trpv4*^+/+^ and *Trpv4*^−/−^ mice. Error bars correspond to s.e.m. (**C**) Evaluation of the Evans blue dye content in retinas from control and STZ *Trpv4*^+/+^ and *Trpv4*^−/−^ mice. Values are mean ± s.e.m. (n = 9-12 per group); *, significant differences of P < 0.05; n.s., not significant.

**Figure 3**. Retinal thickness assessment in control and STZ mice that received a single intravitreal injection of TRPV4 antagonist GSK2193874 on the left eye and vehicle (PBS) on the right eye, 24 h before the end of the 4-week STZ treatment. (**A**) Representative images of non-diffusion weighted images and enlargement corresponding to *white boxes*. Approximate locations of vitreous, retina, and choroid-RPE complex are indicated in *vertical black lines*. (**B**) Representative DWI (**C**) Summary of retinal thickness in four separate groups of control and STZ mice treated with vehicle (*white*) and GSK2193874 (*grey*) (μm, mean ± s.e.m.; n = 10-12). (**D**) Schematic representation of orientated transversal section of the retina equally divided into virtual 20° angles, numbered from 1 to 7, from which retinal thickness (*white double-end arrow*) was measured. Per-section differences of retinal thickness are indicated. ON, optic nerve. *, significant differences of P < 0.05; n.s., not significant.

**Figure 4.** Estimated water diffusion in control and STZ mice that received a single intravitreal injection of TRPV4 antagonist GSK2193874 on the left eye and vehicle (PBS) on the right eye, 24 hours before the end of the 4-week STZ treatment. (**A**) Representative images of ADC maps. (**B**) Summary of ADC profiles in four separate groups. Error bars correspond to s.e.m. (**C**) Schematic representation of transversal retinal section from which ADC was measured. Differences in averaged ADC values were as indicated. *, significant difference (P < 0.05); n.s., not significant.

---

**Supplemental Figure 1**. Follow-up of body weight and glucemia in control and STZ *Trpv4*^+/+^ and *Trpv4*^−/−^ mice. *, significant difference (P < 0.05); **, significant difference (P < 0.025); n.s., not significant.

**Supplemental Figure 2**. Representative DWI image, in which retina was transversally and equally divided into virtual 20° angles to measure retinal thickness. A similar procedure was followed to assess ADC values.

**Supplemental Figure 3**. Anatomical MRI illustrating that some *Trpv4*^−/−^ mice (two out of eleven) showed enlarged lens capsule (delimited by *white dashed circles*) compared with *Trpv4*^+/+^ mice.

## References

Akazawa, Y., Yuki, T., Yoshida, H., Sugiyama, Y., Inoue, S., 2013. Activation of TRPV4 strengthens the tight-junction barrier in human epidermal keratinocytes. Skin Pharmacol Physiol 26, 15–21.

Alessandri-Haber, N., Dina, O.A., Joseph, E.K., Reichling, D., Levine, J.D., 2006. A transient receptor potential vanilloid 4-dependent mechanism of hyperalgesia is engaged by concerted action of inflammatory mediators. J Neurosci 26, 3864–3874.

Alessandri-Haber, N., Dina, O.A., Yeh, J.J., Parada, C.A., Reichling, D.B., Levine, J.D., 2004. Transient receptor potential vanilloid 4 is essential in chemotherapy-induced neuropathic pain in the rat. J Neurosci 24, 4444–4452.

Alessandri-Haber, N., Joseph, E., Dina, O.A., Liedtke, W., Levine, J.D., 2005. TRPV4 mediates pain-related behavior induced by mild hypertonic stimuli in the presence of inflammatory mediator. Pain 118, 70–79.

Alessandri-Haber, N., Yeh, J.J., Boyd, A.E., Parada, C.A., Chen, X., Reichling, D.B., Levine, J.D., 2003. Hypotonicity induces TRPV4-mediated nociception in rat. Neuron 39, 497–511.

Arredondo Zamarripa, D., Noguez Imm, R., Bautista Cortes, A.M., Vazquez Ruiz, O., Bernardini, M., Fiorio Pla, A., Gkika, D., Prevarskaya, N., Lopez-Casillas, F., Liedtke, W., Clapp, C., Thebault, S., 2017. Dual contribution of TRPV4 antagonism in the regulatory effect of vasoinhibins on blood-retinal barrier permeability: diabetic milieu makes a difference. Sci Rep 7, 13094.

Arruda, A.P., Hotamisligil, G.S., 2015. Calcium Homeostasis and Organelle Function in the Pathogenesis of Obesity and Diabetes. Cell Metab 22, 381–397.

Balakrishna, S., Song, W., Achanta, S., Doran, S.F., Liu, B., Kaelberer, M.M., Yu, Z., Sui, A., Cheung, M., Leishman, E., Eidam, H.S., Ye, G., Willette, R.N., Thorneloe, K.S., Bradshaw, H.B., Matalon, S., Jordt, S.E., 2014. TRPV4 inhibition counteracts edema and inflammation and improves pulmonary function and oxygen saturation in chemically induced acute lung injury. Am J Physiol Lung Cell Mol Physiol 307, L158–172.

Bellhorn, R.W., 1984. Analysis of animal models of macular edema. Surv Ophthalmol 28 Suppl, 520–524.

Berkowitz, B.A., Bissig, D., Ye, Y., Valsadia, P., Kern, T.S., Roberts, R., 2012. Evidence for diffuse central retinal edema in vivo in diabetic male Sprague Dawley rats. PLoS One 7, e29619.

Berkowitz, B.A., Gradianu, M., Bissig, D., Kern, T.S., Roberts, R., 2009. Retinal ion regulation in a mouse model of diabetic retinopathy: natural history and the effect of Cu/Zn superoxide dismutase overexpression. Invest Ophthalmol Vis Sci 50, 2351–2358.

Berkowitz, B.A., Grady, E.M., Khetarpal, N., Patel, A., Roberts, R., 2015. Oxidative stress and light-evoked responses of the posterior segment in a mouse model of diabetic retinopathy. Invest Ophthalmol Vis Sci 56, 606–615.

Berkowitz, B.A., Luan, H., Gupta, R.R., Pacheco, D., Seidner, A., Roberts, R., Liggett, J., Knoerzer, D.L., Connor, J.R., Du, Y., Kern, T.S., Ito, Y., 2004. Regulation of the early subnormal retinal oxygenation response in experimental diabetes by inducible nitric oxide synthase. Diabetes 53, 173–178.

Cammalleri, M., Dal Monte, M., Locri, F., Lardner, E., Kvanta, A., Rusciano, D., Andre, H., Bagnoli, P., 2017. Efficacy of a Fatty Acids Dietary Supplement in a Polyethylene Glycol-Induced Mouse Model of Retinal Degeneration. Nutrients 9.

Chen, C.K., Hsu, P.Y., Wang, T.M., Miao, Z.F., Lin, R.T., Juo, S.H., 2018. TRPV4 Activation Contributes Functional Recovery from Ischemic Stroke via Angiogenesis and Neurogenesis. Mol Neurobiol 55, 4127–4135.

Cheung, M., Bao, W., Behm, D.J., Brooks, C.A., Bury, M.J., Dowdell, S.E., Eidam, H.S., Fox, R.M., Goodman, K.B., Holt, D.A., Lee, D., Roethke, T.J., Willette, R.N., Xu, X., Ye, G., Thorneloe, K.S., 2017. Discovery of GSK2193874: An Orally Active, Potent, and Selective Blocker of Transient Receptor Potential Vanilloid 4. ACS Med Chem Lett 8, 549–554.

Clermont, A.C., Aiello, L.P., Mori, F., Aiello, L.M., Bursell, S.E., 1997. Vascular endothelial growth factor and severity of nonproliferative diabetic retinopathy mediate retinal hemodynamics in vivo: a potential role for vascular endothelial growth factor in the progression of nonproliferative diabetic retinopathy. Am J Ophthalmol 124, 433–446.

Cohen, G., Riahi, Y., Sunda, V., Deplano, S., Chatgilialoglu, C., Ferreri, C., Kaiser, N., Sasson, S., 2013. Signaling properties of 4-hydroxyalkenals formed by lipid peroxidation in diabetes. Free Radic Biol Med 65, 978–987.

D’Arceuil, H.E., Westmoreland, S., de Crespigny, A.J., 2007. An approach to high resolution diffusion tensor imaging in fixed primate brain. Neuroimage 35, 553–565.

Daruich, A., Matet, A., Moulin, A., Kowalczuk, L., Nicolas, M., Sellam, A., Rothschild, P.R., Omri, S., Gelize, E., Jonet, L., Delaunay, K., De Kozak, Y., Berdugo, M., Zhao, M., Crisanti, P., Behar-Cohen, F., 2018. Mechanisms of macular edema: Beyond the surface. Prog Retin Eye Res 63, 20–68.

Dysli, C., Enzmann, V., Sznitman, R., Zinkernagel, M.S., 2015. Quantitative Analysis of Mouse Retinal Layers Using Automated Segmentation of Spectral Domain Optical Coherence Tomography Images. Transl Vis Sci Technol 4, 9.

Earley, S., Pauyo, T., Drapp, R., Tavares, M.J., Liedtke, W., Brayden, J.E., 2009. TRPV4-dependent dilation of peripheral resistance arteries influences arterial pressure. Am J Physiol Heart Circ Physiol 297, H1096–1102.

Ebisu, T., Naruse, S., Horikawa, Y., Ueda, S., Tanaka, C., Uto, M., Umeda, M., Higuchi, T., 1993. Discrimination between different types of white matter edema with diffusion-weighted MR imaging. J Magn Reson Imaging 3, 863–868.

Elms, S.C., Toque, H.A., Rojas, M., Xu, Z., Caldwell, R.W., Caldwell, R.B., 2013. The role of arginase I in diabetes-induced retinal vascular dysfunction in mouse and rat models of diabetes. Diabetologia 56, 654–662.

Ganapathy, P.S., Roon, P., Moister, T.K., Mysona, B., Smith, S.B., 2009. Diabetes Accelerates Retinal Neuronal Cell Death In A Mouse Model of Endogenous Hyperhomocysteinemia. Ophthalmol Eye Dis 1, 3–11.

Gao, F., Wang, D.H., 2010. Hypotension induced by activation of the transient receptor potential vanilloid 4 channels: role of Ca2+-activated K+ channels and sensory nerves. J Hypertens 28, 102–110.

Gilliam, J.C., Wensel, T.G., 2011. TRP channel gene expression in the mouse retina. Vision Res 51, 2440–2452.

Grunwald, J.E., DuPont, J., Riva, C.E., 1996. Retinal haemodynamics in patients with early diabetes mellitus. Br J Ophthalmol 80, 327–331.

Hamanaka, K., Jian, M.Y., Weber, D.S., Alvarez, D.F., Townsley, M.I., Al-Mehdi, A.B., King, J.A., Liedtke, W., Parker, J.C., 2007. TRPV4 initiates the acute calcium-dependent permeability increase during ventilator-induced lung injury in isolated mouse lungs. Am J Physiol Lung Cell Mol Physiol 293, L923–932.

Hammes, H.P., Lin, J., Renner, O., Shani, M., Lundqvist, A., Betsholtz, C., Brownlee, M., Deutsch, U., 2002. Pericytes and the pathogenesis of diabetic retinopathy. Diabetes 51, 3107–3112.

Han, B.G., Hao, C.M., Tchekneva, E.E., Wang, Y.Y., Lee, C.A., Ebrahim, B., Harris, R.C., Kern, T.S., Wasserman, D.H., Breyer, M.D., Qi, Z., 2008. Markers of glycemic control in the mouse: comparisons of 6-h- and overnight-fasted blood glucoses to Hb A1c. Am J Physiol Endocrinol Metab 295, E981–986.

Harteneck, C., Reiter, B., 2007. TRP channels activated by extracellular hypo-osmoticity in epithelia. Biochem Soc Trans 35, 91–95.

Hills, C.E., Bland, R., Squires, P.E., 2012. Functional expression of TRPV4 channels in human collecting duct cells: implications for secondary hypertension in diabetic nephropathy. Exp Diabetes Res 2012, 936518.

Hoshi, Y., Okabe, K., Shibasaki, K., Funatsu, T., Matsuki, N., Ikegaya, Y., Koyama, R., 2018. Ischemic Brain Injury Leads to Brain Edema via Hyperthermia-Induced TRPV4 Activation. J Neurosci 38, 5700–5709.

Hossain, A., Heron, D., Davenport, I., Huckaba, T., Graves, R., Mandal, T., Muniruzzaman, S., Wang, S., Bhattacharjee, P.S., 2016. Protective effects of bestatin in the retina of streptozotocin-induced diabetic mice. Exp Eye Res 149, 100–106.

Iuso, A., Krizaj, D., 2016. TRPV4-AQP4 interactions ‘turbocharge’ astroglial sensitivity to small osmotic gradients. Channels (Austin) 10, 172–174.

Jie, P., Tian, Y., Hong, Z., Li, L., Zhou, L., Chen, L., Chen, L., 2015. Blockage of transient receptor potential vanilloid 4 inhibits brain edema in middle cerebral artery occlusion mice. Front Cell Neurosci 9, 141.

Jo, A.O., Ryskamp, D.A., Phuong, T.T., Verkman, A.S., Yarishkin, O., MacAulay, N., Krizaj, D., 2015. TRPV4 and AQP4 Channels Synergistically Regulate Cell Volume and Calcium Homeostasis in Retinal Muller Glia. J Neurosci 35, 13525–13537.

Kaneko, Y., Saito, M., Mori, A., Sakamoto, K., Nakahara, T., Ishii, K., 2006. Vasodilator effects of adrenomedullin on retinal arterioles in streptozotocin-induced diabetic rats. J Ocul Pharmacol Ther 22, 317–322.

Kaplan, H.J., Chiang, C.W., Chen, J., Song, S.K., 2010. Vitreous Volume of the Mouse Measured by Quantitative High-Resolution MRI. ARVO Annual Meeting Abstract 51.

Ke, S.K., Chen, L., Duan, H.B., Tu, Y.R., 2015. Opposing actions of TRPV4 channel activation in the lung vasculature. Respir Physiol Neurobiol 219, 43–50.

Kim, K.J., Ramiro Diaz, J., Iddings, J.A., Filosa, J.A., 2016. Vasculo-Neuronal Coupling: Retrograde Vascular Communication to Brain Neurons. J Neurosci 36, 12624–12639.

Kim, Y.H., Park, S.Y., Park, J., Kim, Y.S., Hwang, E.M., Park, J.Y., Roh, G.S., Kim, H.J., Kang, S.S., Cho, G.J., Choi, W.S., 2012. Reduction of experimental diabetic vascular leakage and pericyte apoptosis in mice by delivery of alphaA-crystallin with a recombinant adenovirus. Diabetologia 55, 2835–2844.

Klaassen, I., Van Noorden, C.J., Schlingemann, R.O., 2013. Molecular basis of the inner blood-retinal barrier and its breakdown in diabetic macular edema and other pathological conditions. Prog Retin Eye Res 34, 19–48.

Kohler, R., Heyken, W.T., Heinau, P., Schubert, R., Si, H., Kacik, M., Busch, C., Grgic, I., Maier, T., Hoyer, J., 2006. Evidence for a functional role of endothelial transient receptor potential V4 in shear stress-induced vasodilatation. Arterioscler Thromb Vasc Biol 26, 1495–1502.

Kohno, T., Ishibashi, T., Inomata, H., Ikui, H., Taniguchi, Y., 1983. Experimental macular edema of commotio retinae: preliminary report. Jpn J Ophthalmol 27, 149–156.

Lakk, M., Yarishkin, O., Baumann, J.M., Iuso, A., Krizaj, D., 2017. Cholesterol regulates polymodal sensory transduction in Muller glia. Glia 65, 2038–2050.

Leal, E.C., Manivannan, A., Hosoya, K., Terasaki, T., Cunha-Vaz, J., Ambrosio, A.F., Forrester, J.V., 2007. Inducible nitric oxide synthase isoform is a key mediator of leukostasis and blood-retinal barrier breakdown in diabetic retinopathy. Invest Ophthalmol Vis Sci 48, 5257–5265.

Li, D., Yang, F., Cheng, H., Liu, C., Sun, M., Wu, K., Ai, M., 2013. Protective effects of total flavonoids from Flos Puerariae on retinal neuronal damage in diabetic mice. Mol Vis 19, 1999–2010.

Li, M.S., Xin, M., Guo, C.L., Lin, G.M., Li, J., Wu, X.G., 2017. Differential expression of breast cancer-resistance protein, lung resistance protein, and multidrug resistance protein 1 in retinas of streptozotocin-induced diabetic mice. Int J Ophthalmol 10, 515–523.

Li, Q., Verma, A., Han, P.Y., Nakagawa, T., Johnson, R.J., Grant, M.B., Campbell-Thompson, M., Jarajapu, Y.P., Lei, B., Hauswirth, W.W., 2010. Diabetic eNOS-knockout mice develop accelerated retinopathy. Invest Ophthalmol Vis Sci 51, 5240–5246.

Liedtke, W., Friedman, J.M., 2003. Abnormal osmotic regulation in trpv4−/− mice. Proc Natl Acad Sci U S A 100, 13698–13703.

Liedtke, W., Kim, C., 2005. Functionality of the TRPV subfamily of TRP ion channels: add mechano-TRP and osmo-TRP to the lexicon! Cell Mol Life Sci 62, 2985–3001.

Liedtke, W., Tobin, D.M., Bargmann, C.I., Friedman, J.M., 2003. Mammalian TRPV4 (VR-OAC) directs behavioral responses to osmotic and mechanical stimuli in Caenorhabditis elegans. Proc Natl Acad Sci U S A 100 Suppl 2, 14531–14536.

Lu, H., Lei, X., 2014. The apparent diffusion coefficient does not reflect cytotoxic edema on the uninjured side after traumatic brain injury. Neural Regen Res 9, 973–977.

Lu, K.T., Huang, T.C., Tsai, Y.H., Yang, Y.L., 2017. Transient receptor potential vanilloid type 4 channels mediate Na-K-Cl-co-transporter-induced brain edema after traumatic brain injury. J Neurochem 140, 718–727.

Martin, P.M., Roon, P., Van Ells, T.K., Ganapathy, V., Smith, S.B., 2004. Death of retinal neurons in streptozotocin-induced diabetic mice. Invest Ophthalmol Vis Sci 45, 3330–3336.

Matsumoto, K., Yamaba, R., Inoue, K., Utsumi, D., Tsukahara, T., Amagase, K., Tominaga, M., Kato, S., 2018. Transient receptor potential vanilloid 4 channel regulates vascular endothelial permeability during colonic inflammation in dextran sulphate sodium-induced murine colitis. Br J Pharmacol 175, 84–99.

Monaghan, K., McNaughten, J., McGahon, M.K., Kelly, C., Kyle, D., Yong, P.H., McGeown, J.G., Curtis, T.M., 2015. Hyperglycemia and Diabetes Downregulate the Functional Expression of TRPV4 Channels in Retinal Microvascular Endothelium. PLoS One 10, e0128359.

Nakazawa, T., Sato, A., Mori, A., Saito, M., Sakamoto, K., Nakahara, T., Ishii, K., 2008. Beta-adrenoceptor-mediated vasodilation of retinal blood vessels is reduced in streptozotocin-induced diabetic rats. Vascul Pharmacol 49, 77–83.

Navaratna, D., McGuire, P.G., Menicucci, G., Das, A., 2007. Proteolytic degradation of VE-cadherin alters the blood-retinal barrier in diabetes. Diabetes 56, 2380–2387.

Nishimura, C., Kuriyama, K., 1985. Alterations in the retinal dopaminergic neuronal system in rats with streptozotocin-induced diabetes. J Neurochem 45, 448–455.

Okada, Y., Shirai, K., Miyajima, M., Reinach, P.S., Yamanaka, O., Sumioka, T., Kokado, M., Tomoyose, K., Saika, S., 2016. Loss of TRPV4 Function Suppresses Inflammatory Fibrosis Induced by Alkali-Burning Mouse Corneas. PLoS One 11, e0167200.

Ozaki, H., Inoue, R., Matsushima, T., Sasahara, M., Hayashi, A., Mori, H., 2018. Serine racemase deletion attenuates neurodegeneration and microvascular damage in diabetic retinopathy. PLoS One 13, e0190864.

Pairet, N., Mang, S., Fois, G., Keck, M., Kuhnbach, M., Gindele, J., Frick, M., Dietl, P., Lamb, D.J., 2018. TRPV4 inhibition attenuates stretch-induced inflammatory cellular responses and lung barrier dysfunction during mechanical ventilation. PLoS One 13, e0196055.

Pannicke, T., Iandiev, I., Wurm, A., Uckermann, O., vom Hagen, F., Reichenbach, A., Wiedemann, P., Hammes, H.P., Bringmann, A., 2006. Diabetes alters osmotic swelling characteristics and membrane conductance of glial cells in rat retina. Diabetes 55, 633–639.

Phuong, T.T.T., Redmon, S.N., Yarishkin, O., Winter, J.M., Li, D.Y., Krizaj, D., 2017. Calcium influx through TRPV4 channels modulates the adherens contacts between retinal microvascular endothelial cells. J Physiol 595, 6869–6885.

Qian, X., Lin, L., Zong, Y., Yuan, Y., Dong, Y., Fu, Y., Shao, W., Li, Y., Gao, Q., 2018. Shifts in renin-angiotensin system components, angiogenesis, and oxidative stress-related protein expression in the lamina cribrosa region of streptozotocin-induced diabetic mice. Graefes Arch Clin Exp Ophthalmol 256, 525–534.

Quiram, P.A., Gonzales, C.R., Schwartz, S.D., 2006. Severe steroid-induced glaucoma following intravitreal injection of triamcinolone acetonide. Am J Ophthalmol 141, 580–582.

Redmon, S.N., Shibasaki, K., Križaj, D., 2017. Transient receptor potential cation channel subfamily V member 4 (TRPV4). Encyclopedia of Signaling Molecules. ed. Choi S, Springer Nature, New York, 1–11.

Reiter, B., Kraft, R., Gunzel, D., Zeissig, S., Schulzke, J.D., Fromm, M., Harteneck, C., 2006. TRPV4-mediated regulation of epithelial permeability. FASEB J 20, 1802–1812.

Ren, X., Li, C., Liu, J., Zhang, C., Fu, Y., Wang, N., Ma, H., Lu, H., Kong, H., Kong, L., 2017. Thioredoxin plays a key role in retinal neuropathy prior to endothelial damage in diabetic mice. Oncotarget 8, 61350–61364.

Roberts, R., Zhang, W., Ito, Y., Berkowitz, B.A., 2003. Spatial pattern and temporal evolution of retinal oxygenation response in oxygen-induced retinopathy. Invest Ophthalmol Vis Sci 44, 5315–5320.

Romero-Aroca, P., 2010. Targeting the pathophysiology of diabetic macular edema. Diabetes Care 33, 2484–2485.

Romero-Aroca, P., Baget-Bernaldiz, M., Pareja-Rios, A., Lopez-Galvez, M., Navarro-Gil, R., Verges, R., 2016. Diabetic Macular Edema Pathophysiology: Vasogenic versus Inflammatory. J Diabetes Res 2016, 2156273.

Ryskamp, D.A., Jo, A.O., Frye, A.M., Vazquez-Chona, F., MacAulay, N., Thoreson, W.B., Krizaj, D., 2014. Swelling and eicosanoid metabolites differentially gate TRPV4 channels in retinal neurons and glia. J Neurosci 34, 15689–15700.

Ryskamp, D.A., Witkovsky, P., Barabas, P., Huang, W., Koehler, C., Akimov, N.P., Lee, S.H., Chauhan, S., Xing, W., Renteria, R.C., Liedtke, W., Krizaj, D., 2011. The polymodal ion channel transient receptor potential vanilloid 4 modulates calcium flux, spiking rate, and apoptosis of mouse retinal ganglion cells. J Neurosci 31, 7089–7101.

Sachdeva, R., Schlotterer, A., Schumacher, D., Matka, C., Mathar, I., Dietrich, N., Medert, R., Kriebs, U., Lin, J., Nawroth, P., Birnbaumer, L., Fleming, T., Hammes, H.P., Freichel, M., 2018. TRPC proteins contribute to development of diabetic retinopathy and regulate glyoxalase 1 activity and methylglyoxal accumulation. Mol Metab 9, 156–167.

Sakamoto, T., Ishibashi, T., Ogura, Y., Shiraga, F., Takeuchi, S., Yamashita, H., Japanese, R., Vitreous Society Triamcinolone Survey, G., 2011. [Survey of triamcinolone-related non-infectious endophthalmitis]. Nippon Ganka Gakkai Zasshi 115, 523–528.

Sood, R., Yang, Y., Taheri, S., Candelario-Jalil, E., Estrada, E.Y., Walker, E.J., Thompson, J., Rosenberg, G.A., 2009. Increased apparent diffusion coefficients on MRI linked with matrix metalloproteinases and edema in white matter after bilateral carotid artery occlusion in rats. J Cereb Blood Flow Metab 29, 308–316.

Stefansson, E., Wilson, C.A., Lightman, S.L., Kuwabara, T., Palestine, A.G., Wagner, H.G., 1987. Quantitative measurements of retinal edema by specific gravity determinations. Invest Ophthalmol Vis Sci 28, 1281–1289.

Suresh, K., Servinsky, L., Reyes, J., Baksh, S., Undem, C., Caterina, M., Pearse, D.B., Shimoda, L.A., 2015. Hydrogen peroxide-induced calcium influx in lung microvascular endothelial cells involves TRPV4. Am J Physiol Lung Cell Mol Physiol 309, L1467–1477.

Thorneloe, K.S., Cheung, M., Bao, W., Alsaid, H., Lenhard, S., Jian, M.Y., Costell, M., Maniscalco-Hauk, K., Krawiec, J.A., Olzinski, A., Gordon, E., Lozinskaya, I., Elefante, L., Qin, P., Matasic, D.S., James, C., Tunstead, J., Donovan, B., Kallal, L., Waszkiewicz, A., Vaidya, K., Davenport, E.A., Larkin, J., Burgert, M., Casillas, L.N., Marquis, R.W., Ye, G., Eidam, H.S., Goodman, K.B., Toomey, J.R., Roethke, T.J., Jucker, B.M., Schnackenberg, C.G., Townsley, M.I., Lepore, J.J., Willette, R.N., 2012. An orally active TRPV4 channel blocker prevents and resolves pulmonary edema induced by heart failure. Sci Transl Med 4, 159ra148.

Vergnolle, N., Cenac, N., Altier, C., Cellars, L., Chapman, K., Zamponi, G.W., Materazzi, S., Nassini, R., Liedtke, W., Cattaruzza, F., Grady, E.F., Geppetti, P., Bunnett, N.W., 2010. A role for transient receptor potential vanilloid 4 in tonicity-induced neurogenic inflammation. Br J Pharmacol 159, 1161–1173.

Villalta, P.C., Rocic, P., Townsley, M.I., 2014. Role of MMP2 and MMP9 in TRPV4-induced lung injury. Am J Physiol Lung Cell Mol Physiol 307, L652–659.

Vincent, F., Duncton, M.A., 2011. TRPV4 agonists and antagonists. Curr Top Med Chem 11, 2216–2226.

Willermain, F., Scifo, L., Weber, C., Caspers, L., Perret, J., Delporte, C., 2018. Potential Interplay between Hyperosmolarity and Inflammation on Retinal Pigmented Epithelium in Pathogenesis of Diabetic Retinopathy. Int J Mol Sci 19.

Willette, R.N., Bao, W., Nerurkar, S., Yue, T.L., Doe, C.P., Stankus, G., Turner, G.H., Ju, H., Thomas, H., Fishman, C.E., Sulpizio, A., Behm, D.J., Hoffman, S., Lin, Z., Lozinskaya, I., Casillas, L.N., Lin, M., Trout, R.E., Votta, B.J., Thorneloe, K., Lashinger, E.S., Figueroa, D.J., Marquis, R., Xu, X., 2008. Systemic activation of the transient receptor potential vanilloid subtype 4 channel causes endothelial failure and circulatory collapse: Part 2. J Pharmacol Exp Ther 326, 443–452.

Xu, H., Fu, Y., Tian, W., Cohen, D.M., 2006. Glycosylation of the osmoresponsive transient receptor potential channel TRPV4 on Asn-651 influences membrane trafficking. Am J Physiol Renal Physiol 290, F1103–1109.

Yang, X.F., Huang, Y.X., Lan, M., Zhang, T.R., Zhou, J., 2018. Protective Effects of Leukemia Inhibitory Factor on Retinal Vasculature and Cells in Streptozotocin-induced Diabetic Mice. Chin Med J (Engl) 131, 75–81.

Yanoff, M., Fine, B.S., Brucker, A.J., Eagle, R.C., Jr., 1984. Pathology of human cystoid macular edema. Surv Ophthalmol 28 Suppl, 505–511.

Ye, L., Kleiner, S., Wu, J., Sah, R., Gupta, R.K., Banks, A.S., Cohen, P., Khandekar, M.J., Bostrom, P., Mepani, R.J., Laznik, D., Kamenecka, T.M., Song, X., Liedtke, W., Mootha, V.K., Puigserver, P., Griffin, P.R., Clapham, D.E., Spiegelman, B.M., 2012. TRPV4 is a regulator of adipose oxidative metabolism, inflammation, and energy homeostasis. Cell 151, 96–110.

Yin, J., Michalick, L., Tang, C., Tabuchi, A., Goldenberg, N., Dan, Q., Awwad, K., Wang, L., Erfinanda, L., Nouailles, G., Witzenrath, M., Vogelzang, A., Lv, L., Lee, W.L., Zhang, H., Rotstein, O., Kapus, A., Szaszi, K., Fleming, I., Liedtke, W.B., Kuppe, H., Kuebler, W.M., 2016. Role of Transient Receptor Potential Vanilloid 4 in Neutrophil Activation and Acute Lung Injury. Am J Respir Cell Mol Biol 54, 370–383.

Yushkevich, P.A., Piven, J., Hazlett, H.C., Smith, R.G., Ho, S., Gee, J.C., Gerig, G., 2006. User-guided 3D active contour segmentation of anatomical structures: significantly improved efficiency and reliability. Neuroimage 31, 1116–1128.

Zhao, H., Zhang, K., Tang, R., Meng, H., Zou, Y., Wu, P., Hu, R., Liu, X., Feng, H., Chen, Y., 2018. TRPV4 Blockade Preserves the Blood-Brain Barrier by Inhibiting Stress Fiber Formation in a Rat Model of Intracerebral Hemorrhage. Front Mol Neurosci 11, 97.

Zhao, P.Y., Gan, G., Peng, S., Wang, S.B., Chen, B., Adelman, R.A., Rizzolo, L.J., 2015. TRP Channels Localize to Subdomains of the Apical Plasma Membrane in Human Fetal Retinal Pigment Epithelium. Invest Ophthalmol Vis Sci 56, 1916–1923.

Zheng, L., Du, Y., Miller, C., Gubitosi-Klug, R.A., Kern, T.S., Ball, S., Berkowitz, B.A., 2007. Critical role of inducible nitric oxide synthase in degeneration of retinal capillaries in mice with streptozotocin-induced diabetes. Diabetologia 50, 1987–1996.

Zhou, K.K., Benyajati, S., Le, Y., Cheng, R., Zhang, W., Ma, J.X., 2014. Interruption of Wnt signaling in Muller cells ameliorates ischemia-induced retinal neovascularization. PLoS One 9, e108454.

